# Patient-derived organoids from malignant pleural effusion to explore for alternative therapies in thoracic tumors

**DOI:** 10.64898/2026.03.04.709498

**Authors:** Rocio Ferreiro-Miguens, Isabel Diez-Grandio, Roi Soto-Feijoo, Lucía Ferreiro, Jorge Garcia, María Otero-Alen, Ihab Abdulkader, Beatriz Bernardez, Eduardo Dominguez, Miguel Abal, Luis Leon-Mateos

## Abstract

Thoracic malignancies, including lung adenocarcinoma (ADC) and malignant pleural mesothelioma (MPM), remain associated with poor prognosis and limited durable therapeutic responses in advanced stages. Although targeted therapies and immunotherapy have improved outcomes in selected patients, systemic chemotherapy continues to play a central role in routine clinical practice. However, treatment response is highly heterogeneous, and reliable predictive biomarkers of chemotherapy sensitivity are lacking. Both ADC and MPM frequently involve the pleural cavity and are commonly associated with malignant pleural effusion (MPE), which contributes to symptoms such as dyspnea and chest pain and requires therapeutic drainage. Importantly, MPE represents a clinically accessible source of viable tumor cells obtained through minimally invasive procedures.

In this study, we established patient-derived organoids (PDOs) from malignant pleural effusion samples obtained from five patients with advanced lung adenocarcinoma and, as an exploratory extension, from one patient with malignant pleural mesothelioma. Organoids were characterized by immunohistochemistry and subjected to systematic chemotherapy drug screening. Inter-model variability in treatment response was assessed, and selected drug sensitivities were further validated through dose–response assays.

Pleural effusion–derived organoids successfully recapitulated tumor-specific phenotypic features and revealed marked heterogeneity in chemotherapy sensitivity across models. Secondary validation confirmed the reproducibility of selected responses.

Our findings support the feasibility of generating functional organoid models from malignant pleural effusions and highlight their potential as translational platforms for individualized chemotherapy profiling in advanced thoracic malignancies.

## INTRODUCTION

Thoracic malignancies represent a major cause of cancer-related morbidity and mortality worldwide. Among them, non-small cell lung cancer (NSCLC) accounts for approximately 85% of all cases, with lung adenocarcinoma (ADC) being its most common histological subtype (1,2). Despite significant therapeutic advances, including targeted therapies and immune checkpoint inhibitors, a substantial proportion of patients either lack actionable molecular alterations or eventually develop resistance to treatment (3–5). As a result, systemic chemotherapy continues to play a central role in the management of advanced disease (6). Malignant pleural mesothelioma (MPM) is a distinct but similarly aggressive thoracic malignancy arising from the pleural mesothelium (7,8).Although less frequent than lung cancer, mesothelioma is characterized by poor prognosis and limited durable therapeutic benefit. Platinum-based chemotherapy remains a cornerstone of systemic treatment, with modest survival improvements (9). In both ADC and MPM, the management of advanced disease remains challenging: chemotherapy responses are heterogeneous, durable benefit is limited in many patients, reliable functional predictors of treatment efficacy are lacking, and treatment decisions are frequently empirical, highlighting an important unmet clinical need (10,11).

Beyond their biological differences, lung adenocarcinoma and mesothelioma share important clinical features, particularly in advanced stages. Both malignancies commonly involve the pleural cavity and frequently lead to the development of malignant pleural effusion (MPE) (12,13). The accumulation of pleural fluid contributes to symptoms such as progressive dyspnea, chest pain, cough, and reduced functional status, significantly impairing quality of life (14). Therapeutic thoracentesis or pleural drainage is often required for symptomatic relief and diagnostic confirmation. Thus, malignant pleural effusion represents not only a hallmark of advanced thoracic disease but also a routinely accessible source of viable tumor cells obtained through minimally invasive procedures. In this context, MPE represent a minimally invasive source of cancer cells that may better capture tumor heterogeneity, with limited stromal cell contamination (15), and offer a unique opportunity to obtain patient-specific tumor material at a clinically relevant time point, often when therapeutic decisions are urgently needed.

To this regard, current preclinical thoracic models including two- and three-dimensional in vitro models and genetically engineered murine models, have shown limited capacity to recapitulate these diseases, and there is general consensus concluding that high quality preclinical models are essential for the development of new treatments for this lethal cancer (16). Patient-derived organoids (PDOs) have recently emerged as promising three-dimensional ex vivo models capable of preserving tumor architecture, cellular heterogeneity, and phenotypic characteristics of the original malignancy (17,18). Compared with conventional two-dimensional cell cultures, organoids provide improved physiological relevance for drug response assessment and have demonstrated utility in multiple solid tumors, including lung cancer with PDOs mainly derived from surgical specimens but also from malignant pleural effusion with favorable success (19). Preliminary reports also support the feasibility of generating organoid cultures from mesothelioma specimens (20). However, the systematic use of pleural effusion–derived organoids for functional chemotherapy profiling in thoracic malignancies remains insufficiently explored.

In this study, we established patient-derived organoids from malignant pleural effusion samples obtained from patients with advanced lung adenocarcinoma and, as an exploratory extension, from a case of malignant pleural mesothelioma. We applied a standardized workflow to characterize these models and to functionally evaluate their response to systemic chemotherapy. Specifically, we aimed to (i) generate and validate pleural effusion–derived organoids, (ii) assess inter-patient heterogeneity in chemotherapy sensitivity, and (iii) confirm selected drug responses through secondary validation assays. By integrating clinically accessible tumor material with functional drug testing, we sought to explore the translational potential of organoid-based profiling as a complementary strategy to current therapeutic decision-making in advanced thoracic malignancies.

## METHODS

### Sample collection

Malignant pleural effusion (MPE) samples were obtained from patients diagnosed with advanced thoracic malignancies, including five cases of lung adenocarcinoma (ADC) and one case of malignant pleural mesothelioma (MPM). All patients presented with symptomatic pleural effusion requiring diagnostic or therapeutic drainage. Clinical diagnosis was established by histopathological examination and immunohistochemical analysis of pleural biopsy or cytological specimens.

The study was approved by the Galician Research Ethics Committee (reference number 2017/538) and written informed consent was obtained from all participating patients before their enrolment in the study.

Approximately 100 mL of pleural fluid per patient was collected under sterile conditions. Samples were processed within 2 hours of collection. Pleural fluid was centrifuged at 500 × g for 10 minutes at 4°C to obtain a cellular pellet. The pellet was washed with cold phosphate-buffered saline (PBS) and treated with red blood cell lysis buffer for 10 minutes at 37°C to eliminate erythrocytes. Cells were subsequently washed with cold PBS and centrifuged. Following processing, the resulting cellular suspension was divided into two fractions: one aliquot was cryopreserved in fetal bovine serum (FBS) containing 10% dimethyl sulfoxide (DMSO), while the remaining viable cells were used for immediate establishment of three-dimensional organoid cultures.

### PDO culture and expansion

The cells were embedded in 40 µL droplets of a three-dimensional matrix composed of 70% basement membrane extract (BME) and 30% organoid culture medium. After polymerization, culture medium was added. The organoid culture medium consisted of DMEM/F-12 supplemented with epidermal growth factor (EGF), fibroblast growth factor 10 (FGF10), and additional niche-supporting additives to sustain thoracic tumor growth. Cultures were maintained at 37°C in a humidified incubator with 5% CO₂, and the medium was refreshed every 2–3 days. Three-dimensional structures typically emerged within 7–10 days after seeding, progressively developing into compact, irregularly shaped spheroids. Once established, organoids were expanded through mechanical and/or enzymatic dissociation and re-embedded in fresh BME for subsequent passages and downstream applications.

### Immunohistochemistry

Organoids were fixed in 4% paraformaldehyde (PFA), embedded in agarose, and subsequently processed for paraffin embedding. Sections were stained with hematoxylin and eosin (H&E) for morphological evaluation.

Immunohistochemical analysis was performed using markers selected according to the original tumor diagnosis. For lung adenocarcinoma derived organoids, the panel included Thyroid Transcription Factor 1 (TTF-1), cytokeratin 7 (CK7), and Napsin A, when applicable. For malignant pleural mesothelioma derived organoids, markers included calretinin, Wilms Tumor 1 (WT-1), and BRCA1-associated protein 1 (BAP1). Epithelial markers such as MOC-31 and Ber-EP4 were used when necessary to support differential diagnosis.

### Pharmacological study

Patient-derived organoids were cultured for 1–2 weeks, dissociated into single cells, resuspended in PDO medium with BME, and seeded into 384-well plates. A total of 169 compounds were screened using the Echo 650 Liquid Handler, and LPDOs were incubated with drugs at 10 μM for 72 hours under standard culture conditions. Cell viability was measured with the alamar-blue reagent for the fluorimetric quantification of resazurin reduction by living cells. Data were analyzed with GraphPad Prism (v.10.1.1). To assess the translational potential of this PDO model, we used the NCI Approved Oncology Set (169 compounds), comprising FDA-approved agents with clinical activity across solid and hematologic malignancies. The library includes conventional chemotherapies and targeted therapies, such as tyrosine kinase inhibitors, providing a broad assessment of drug sensitivities that may guide therapeutic strategies in adenocarcinoma.

Cell viability assay metric parameters were evaluated in all screens run (7 PDO), averaging Źfactor < 0.5, Signal to Background (S/B) = 3.5, and Coefficient of Variation (%) = 15.2, confirming the feasibility of our method to distinguish candidates influencing the viability of PDO. The threshold to identify hits for IC50 quantitation studies was established at 90% of inhibition.

Dose-response curves were performed by adding 8 concentrations of 1:3 serial dilutions of 10 μM top dose by using the Echo 650 Liquid Handler. Data were analyzed by fitting signals to the nonlinear statistical analysis Variable Slope model (4-parameter logistic equation) using the GraphPad Prism (v.10.1.1) program.

## RESULTS

### Establishment and Immunophenotypic Characterization of Pleural Effusion–Derived NSCLC Organoids

Five patients diagnosed non-small cell lung cancer (NSCLC) were included in the study, the swimmer plot in Figure 1A illustrating the clinical histories including sequential chemotherapy regimens and treatment modifications over time, together with the clinical characteristics of each patient. The median age at diagnosis was 64.6 years (range, 48–87), and the majority of patients were male. Most NSCLC cases were diagnosed at stage IV disease. None of the NSCLC patients harbored actionable molecular alterations; two cases presented no detectable targetable mutations, while other two patients exhibited TP53 mutations without available targeted therapeutic options. These findings corresponded to patients with limited availability of precision therapies and the consequent reliance on systemic chemotherapy.

**Figure 1.**
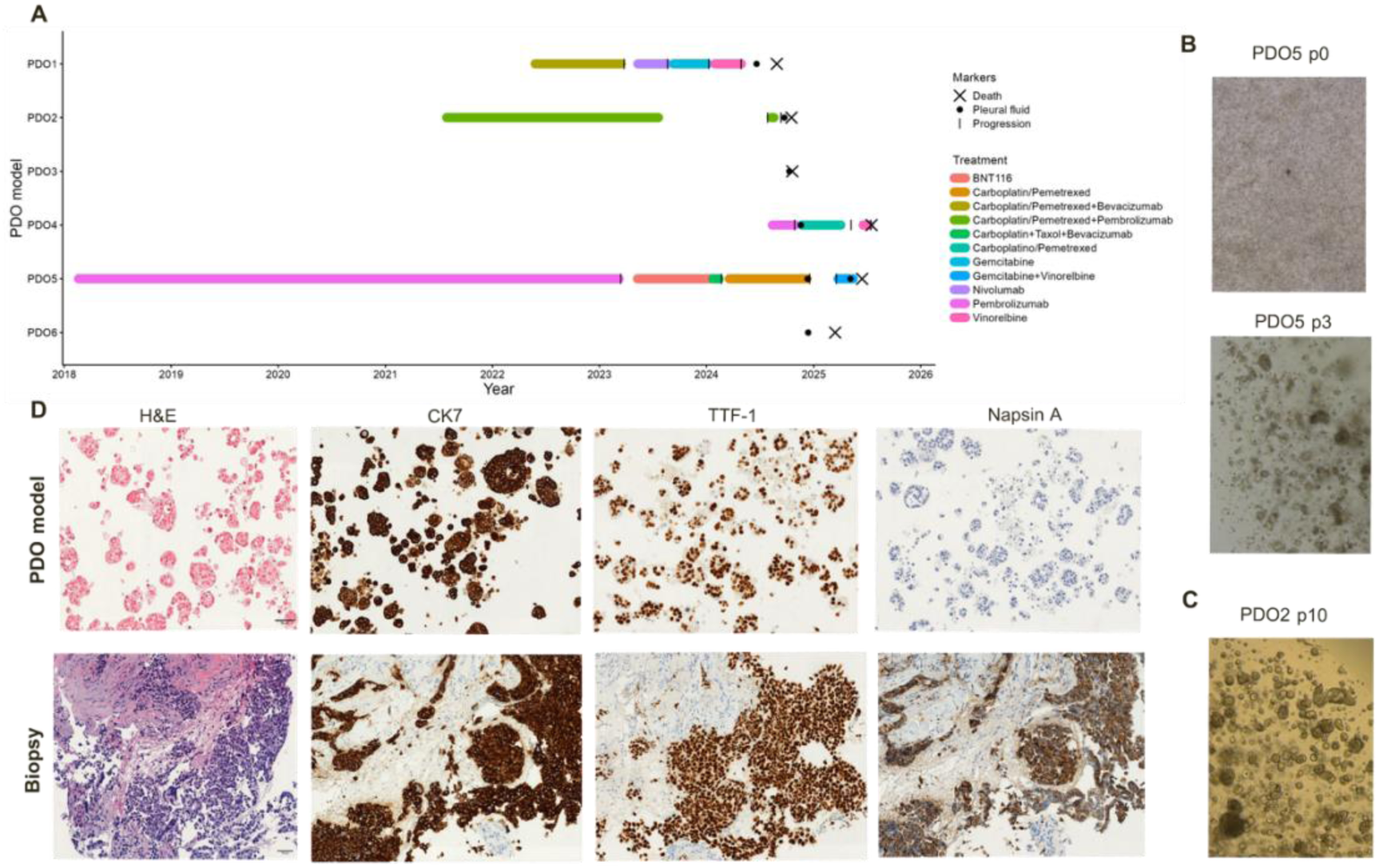
**Clinical course, organoid establishment, and histologic characterization of pleural effusion–derived NSCLC organoids**. (A) Swimmer plot showing the clinical timeline of NSCLC patients, including diagnosis, systemic treatments, development of malignant pleural effusion (MPE), pleural fluid collection for organoid generation, and survival status. (B) Representative brightfield images of PDO5 at early (p0) and later (p3) passages. (C) Representative brightfield image of PDO2 at passage 10 showing compact three-dimensional organoid growth. (D) Histologic and immunohistochemical comparison between a representative NSCLC-PDO (PDO4, upper row) and the corresponding primary bronchial biopsy (lower row), including H&E, CK7, TTF-1, and Napsin A staining.

Malignant pleural effusion (MPE) developed at variable time points during the disease course. In several cases, MPE occurred after multiple lines of systemic treatment, reflecting advanced and heavily pretreated disease. In contrast, in two cases (PDO3 and PDO6), pleural effusion was present at baseline, representing an early manifestation of advanced-stage malignancy. The time from diagnosis to MPE ranged from 97 to 2487 days among evaluable cases. Pleural fluid collection for organoid generation is indicated for each patient (Figure 1A), and occurred in the context of active disease, frequently during or after progression on prior therapy. All patients ultimately succumbed to their disease, with overall survival ranging from 10 to 2,676 days. Together, these data underscore the aggressive clinical course and limited therapeutic options in this cohort, particularly in the absence of actionable molecular alterations, providing a clinically relevant context for functional drug profiling using pleural effusion–derived organoids (Table 1).

**Table 1.**
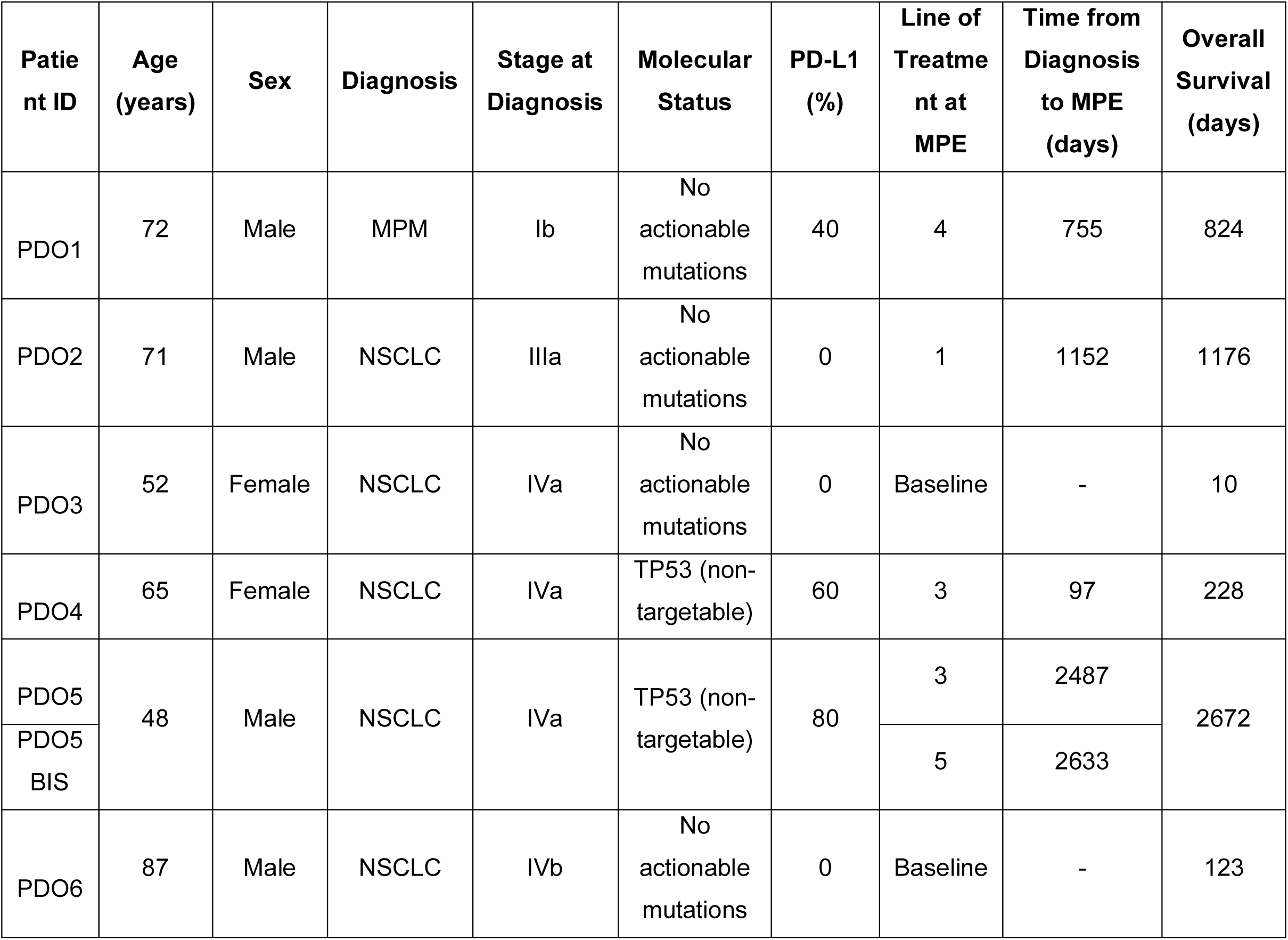
Clinical and molecular characteristics of patients included in the study. Demographic, clinicopathological, and molecular features of patients from whom pleural effusion–derived organoids were generated. The table summarizes age, sex, diagnosis, stage at diagnosis, molecular status, PD-L1 tumor proportion score (TPS), line of treatment at the time of malignant pleural effusion (MPE), time from diagnosis to MPE (days), and overall survival (days). PDO5 and PDO5 BIS correspond to longitudinal pleural effusion samples obtained from the same patient at different timepoints during disease progression.

The generation of the patient-derived organoids from malignant pleural effusion was performed within the two hours upon drainage by ultrasound-guided thoracentesis. Approximately, 100 cc of malignant pleural fluid was processed at 4°C as described Arias-Diaz et al., 2023 (21), for cellular enrichment and removal of erythrocytes through red cell blood lysis. The resulting cellular pellet containing the NSCLC tumoral compartment, as well as the microenvironment, was embedded in Basement Membrane Extract (BME), that contains laminin, collagen IV, entactin, and heparan sulfate proteoglycans, and is specifically designed to support the establishment and expansion of robust organoid cultures by mimicking the in vivo microenvironment. Upon BME gelation, the purified NSCLC cells were cultured in the presence of growth-factor enriched organoid medium composed of DMEM/F12 supported by a cocktail of antioxidants, fatty acids, proteins, small molecules as a ROCK inhibitor, and growth and signaling factors containing epithelial-cell mitogens, such as epidermal growth factor (EGF) and fibroblast growth factor 10 (FGF10), to maintain epithelial-cell character.

Organoid’s formation was evidenced at weeks 0-2 depending on the NSCLC-PDO models, displaying a heterogeneous cellular composition, including epithelial tumor cells and microenvironment-associated elements. During the initial passages the tumoral compartment was progressively enriched while the microenvironment compartment disappeared (Figure 1B). The successful rate of NSCLC-PDO model generation from pleural effusion was 70%, with the observed dense and compact morphology of the organoids indicative of a structural cellular arrangement reminiscent of that of a NSCLC tumor (Figure 1C).

Histologic comparison between the primary bronchial biopsy and the corresponding patient-derived organoid (PDO4) was performed using defined morphologic and immunophenotypic criteria (Figure 1D and Supplementary Figure 1). Hematoxylin and eosin (H&E) staining of the primary tumor revealed invasive acinar-pattern adenocarcinoma characterized by irregular glandular structures embedded in desmoplastic stroma. Tumor cells exhibited enlarged hyperchromatic nuclei, moderate nuclear pleomorphism, prominent nucleoli, increased nuclear-to-cytoplasmic ratio, and partial loss of epithelial polarity (Figure 1D, lower row). In parallel, the PDO demonstrated well-formed three-dimensional glandular structures with clear luminal differentiation, closely resembling the acinar growth pattern observed in the original tumor (Figure 1D, upper row). Cytologically, the organoid retained enlarged nuclei, conspicuous nucleoli, increased nuclear-to-cytoplasmic ratio, moderate cytoplasmic volume, and partial epithelial polarity, with cohesive cellular organization. The absence of desmoplastic stroma in the PDO is consistent with the in vitro three-dimensional culture system. Immunohistochemically, both the primary tumor and the PDO showed diffuse cytoplasmic CK7 expression and strong nuclear TTF-1 positivity, confirming preservation of pulmonary adenocarcinoma differentiation. While Napsin A expression was diffusely positive in the primary tumor, it was not detected in the PDO. Overall, the organoid faithfully recapitulated the principal histomorphologic and lineage-defining immunophenotypic characteristics of the patient’s lung adenocarcinoma (Figure 1D; comparable epithelial differentiation and malignant cytological features were observed across the remaining NSCLC-derived organoid models, Supplementary Figure 1). Collectively, these findings demonstrate the feasibility of generating robust and reliable NSCLC patient-derived organoids directly from malignant pleural effusions, supporting their use as preclinical models representative of advanced-stage disease.

### Drug screening

Once we established this representative set of NSCLC-PDO models, we designed a high throughput screening strategy for the identification of alternative therapies that may improve the outcome of this deadly disease. For this, the fluorometric AlamarBlue cell viability assay detecting cellular metabolic activity by conversion of the blue resazurin dye to the highly fluorescent red resorufin, has been miniaturized and scaled to a 384-well culture plate to screen the inhibitory effects of FDA-approved anticancer agents using the Approved Oncology Set (version X) provided by the Developmental Therapeutics Program (DTP) of the National Cancer Institute (NCI), NIH. This annotated drug library comprises 169 anticancer agents, each with documented clinical activity across various solid and hematologic malignancies. The set is designed to cover broad pharmacological mechanisms, enabling comprehensive interrogation of cancer-specific dependencies and drug sensitivity profiles. The compounds span a wide range of mechanistic classes, including cytotoxic chemotherapies (e.g., taxanes, platinum salts, anthracyclines, antimetabolites), DNA-damaging agents and topoisomerase inhibitors, as well as targeted therapies, such as tyrosine kinase inhibitors, PARP inhibitors, CDK4/6 inhibitors, and epigenetic modulators. These agents collectively cover the major hallmarks of cancer, including cell cycle regulation, DNA damage repair, apoptosis evasion, angiogenesis, immune evasion, and metabolic reprogramming. Their inclusion in this set reflects prior clinical efficacy evidence and pharmacokinetic profiles compatible with in vitro screening settings. Organoids culture was incubated in the presence of 10 µM concentration of each of the FDA-approved anticancer drugs for 72 hours before revealed with AlamarBlue assay. A representative image of the HTS 384-well plate from PDO4 model is shown in Figure 2A, with the wells with organoids colored in magenta to blue indicative of the cellular viability to cell toxicity, respectively, and the panels on the right showing a magnification of representative examples of viable untreated organoids (upper panel), control of cell toxicity with carboplatin at 500 µM (middle panel), and an active compound with massive apoptotic effect (lower panel).

**Figure 2.**
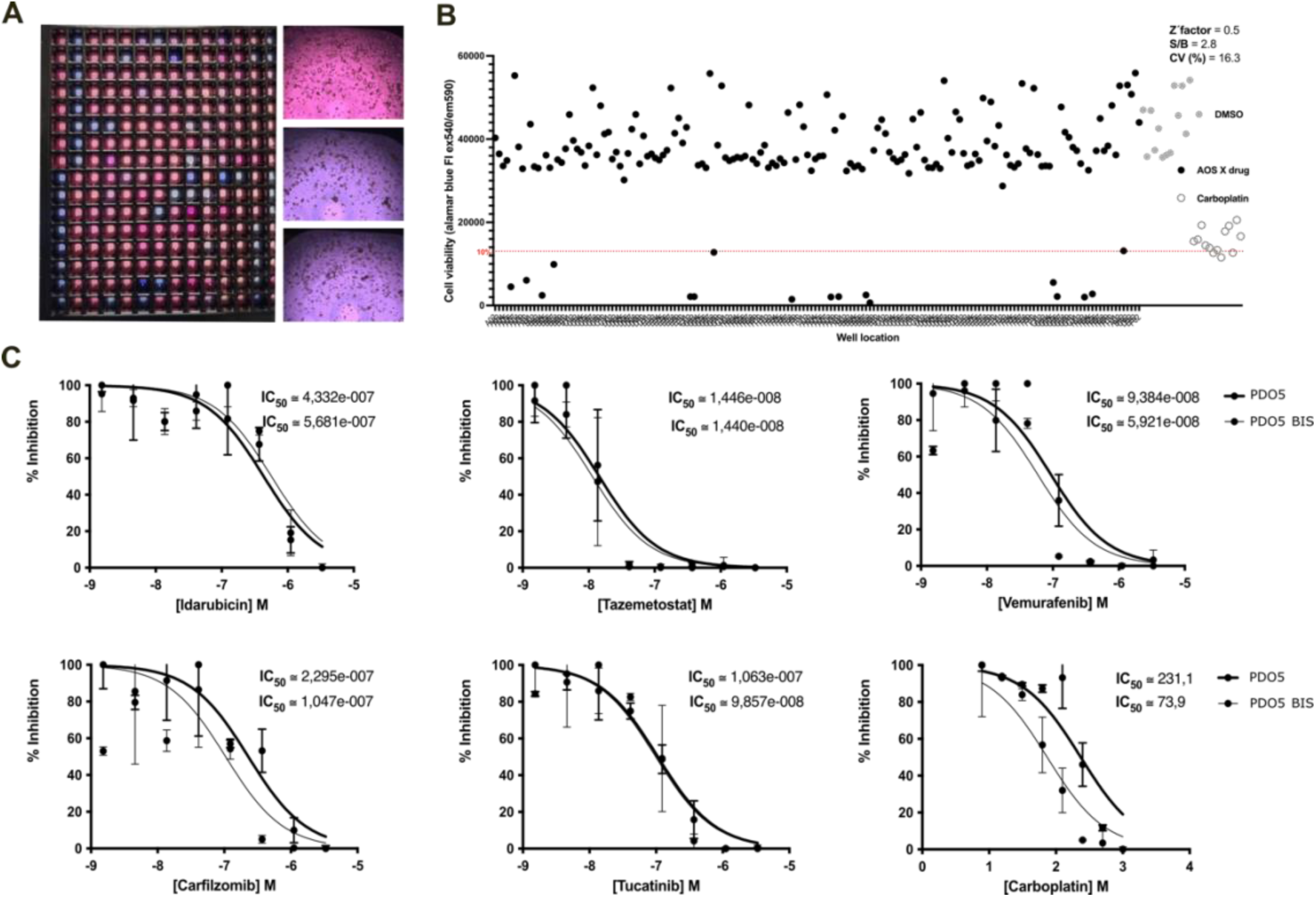
High-throughput drug screening and dose–response validation in pleural effusion–derived NSCLC organoids. (A) Representative image of the 384-well high-throughput screening plate for PDO4 treated with the NCI Approved Oncology Set (169 FDA-approved anticancer agents) at 10 µM for 72 hours. Wells are color-coded from magenta (high viability) to blue (low viability). Panels on the right show representative images of untreated organoids (upper panel), carboplatin-treated control at 500 µM (middle panel), and an active compound inducing marked cytotoxicity (lower panel). (B) Representative scatter plot of PDO4 screening showing drug-induced viability measured by AlamarBlue fluorescence (540/590 nm). DMSO-treated wells define maximal proliferation, and 500 µM carboplatin represents the toxicity control; scatter plots from the remaining NSCLC-PDO models are shown in Supplementary Figure 2. Compounds with marked reduction in viability were considered hits. (C) Dose–response validation of selected common hits in longitudinal paired models (PDO5 and PDO5 BIS). Percent inhibition is plotted against drug concentration (log M). IC₅₀ values are indicated for idarubicin, tazemetostat, vemurafenib, carfilzomib, tucatinib, and carboplatin.

Figure 2B illustrates the representative scatter plot of PDO4 screening with the 169 FDA-approved anticancer agents (black dots) distributed as per cell viability based on Alamarblue fluorescent signal (540/590nm), with the window of viability defined by the DSMO control as maximal NSCLC-PDO proliferation (grey dots) and the 500 µM Carboplatin as control of toxicity representing 10% of organoids viability (white dots; Supplementary Figure 2). Those compounds showing higher toxicity when exposed to the NSCLC-PDOs were considered as hits. In addition, those common hits identified in at least 4 of the NSCLC-PDO models were selected for further validation in a dose-response cell viability assay. These common hits included the competitive kinase inhibitor Vemurafenib, with activity against BRAF kinase with mutations like V600E; Tucatinib, a tyrosine kinase inhibitor that targets the human epidermal growth factor receptor 2 (HER2); the proteasome inhibitor Carfilzomib; or the anthracycline Idarubicin, that intercalate into DNA and also inhibits Topoisomerase II activity. Finally, we also identified the specific EZH2 methyltransferase inhibitor Tazemetostat and the histone deacetylase inhibitors Panobinostat and Romidepsin, with cytotoxic activity in these NSCLC-PDO preclinical models.

The dose-response validation was conducted in the PDO5 model (Figure 2C). Moreover, we generated two paired NSCLC-PDO5 models from longitudinal pleural effusions of this particular patient, collected at two different timepoints during the clinical history with six months of interval between the two thoracocentesis, resulting in PDO5 and PDO5 BIS models. The dose-response Alamarblue assay included the following concentrations of each of the seven common hits identified: 3.33E-06, 1.11E-06, 3.70E-07, 1.23E-07, 4.12E-08, 1.37E-08, 4.58E-09 molar. As shown in Figure 2C, Idarubicin (upper left panel; IC50 4,3E-07M in NSCLC-PDO5 and 5,7E-07 in NSCLC-PDO5bis), Tazemetostat (upper middle panel; IC50 1,4E-08M in both NSCLC-PDO5 and NSCLC-PDO5bis), Vemurafenib (upper right panel; IC50 9,4E-08M in NSCLC-PDO5 and 5,9E-08 in NSCLC-PDO5bis), Carfilzomib (lower left panel; IC50 2,3E-07M in NSCLC-PDO5 and 1,0E-07 in NSCLC-PDO5bis), and Tucatinib (lower middle panel; IC50 1,1E-07M in NSCLC-PDO5 and 9,9E-08 in NSCLC-PDO5bis) were validated as active compounds in these NSCL-PDO models, while Panobinostat and Romidepsin did not confirmed the cytotoxic activity shown in the high-throughput screening. Moreover, a dose-response analysis of these paired NSCLC-PDO5 models with Carboplatin showed activity at high concentration (lower right panel; (IC50 231E-03M in NSCLC-PDO5 and 74E-03 in NSCLC-PDO5bis; 135E-03 as maximal tolerated dose in plasma of patients (22)), compatible with a clinical setting of resistant disease. In addition to validate the selected hits as compounds with potential activity in NSCLC, all these results first highlight a robust reproducibility of the paired PDO models from two different longitudinal samples of the patient advanced disease of origin; and second, recapitulate the absence of response of the patient to Carboplatin based therapy to further confirm the reliability of these NSCLC-PDOs as valuable preclinical models to conduct pharmacological studies to design alternative therapeutic strategies.

### Mesothelioma case

We wanted to complement this pharmacological approach in NSCLC-PDOs by presenting the generation of pleural effusion organoids derived from a malignant mesothelioma patient (MPM-PDO; malignant pleural mesothelioma-patient derived organoids), also representing a poor prognosis disease with reduced targeted therapies and limited preclinical models. Briefly, a 72-year-old male with a history of asbestos exposure presented with progressive and persistent dyspnea and non-productive cough. The patient was afebrile and normotensive, with a saturation of 95%, and showed a right pleural effusion by chest X-ray, interpreted as a parapneumonic effusion. A PET-CT scan showed a right pleural effusion with nodular thickening of the pleural layers, possibly related to a metastatic non small cell lung cancer or mesothelioma (Figure 3A; left panels, February 2022). This study was completed with ultrasound-guided thoracentesis, draining a total of 800 cc of pleural effusion, without complications. The cytological analysis of the effusion confirmed the diagnosis of malignant mesothelioma. Immunohistochemistry demonstrated strong nuclear and cytoplasmic positivity for Calretinin and nuclear expression of Wilms Tumor 1 (WT1), together with complete loss of nuclear staining for BRCA1-associated protein 1 (BAP1). In contrast, epithelial markers including TTF-1, MOC-31, and Ber-EP4 were consistently negative, supporting the mesothelial origin of the neoplastic cells and ruling out an adenocarcinomatous process. In addition, the levels of expression of Programmed Death Ligand-1 (PD-L1) tumor proportion score (TPS) was 40%. A multidisciplinary committee ruled out surgery due to the extensive nature of the tumor. Given the history of active ulcerative colitis and the lack of authorization in our country for the first-line combination of Nivolumab and Ipilimumab for the epithelioid subtype, treatment was initiated with a regimen of Carboplatin, Pemetrexed, and Bevacizumab every three weeks. The patient achieved a partial radiological response and clinical improvement, with decreased of dyspnea and cough. Unfortunately, 13 months later, progression of the disease was observed with an increase in the size of the right pleural implants and worsening of pleural effusion (Figure 3A; middle panels, March 2023). At this moment, monotherapy with Nivolumab 360 mg every 3 weeks was started, with no relevant toxicities or clinical changes but lamentably the patient presented a new tumor progression in the first radiological evaluation. The patient received a third line of treatment with Gemcitabine 1000 mg IV at days 1 and 8 every 3 weeks and, upon a new tumoral progression (Figure 3A; right panels, January 2024), NGS analysis was performed on tissue obtained from the pleural biopsy. No driver mutations were found in the molecular study and a fourth line of chemotherapy was indicated with Vinorelbine 60 mg/m2 orally on days 1 and 8 every 3 weeks.

**Figure 3.**
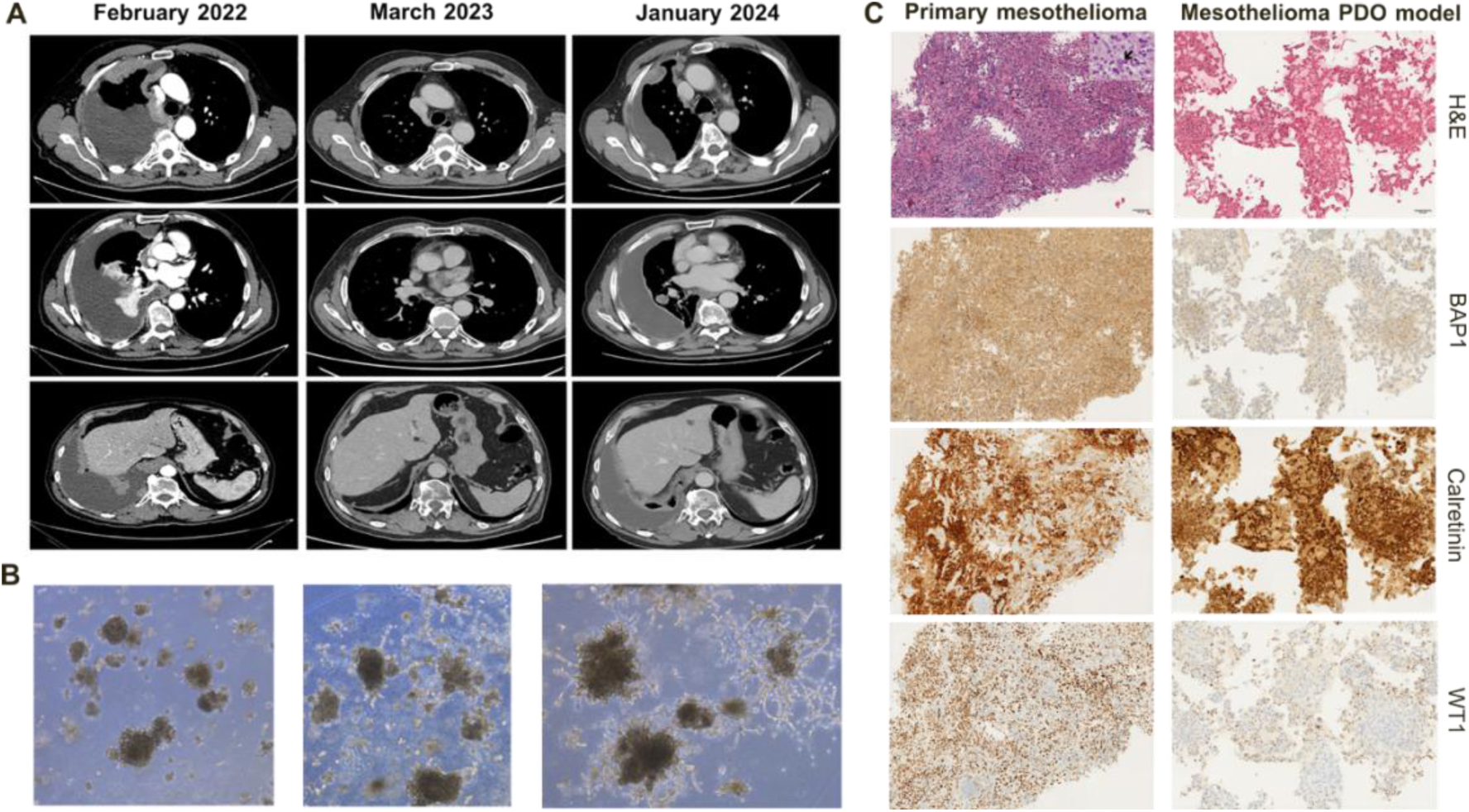
Clinical course and establishment of a pleural effusion–derived mesothelioma organoid model. (A) Serial PET-CT images obtained at diagnosis (February 2022), first progression (March 2023), and subsequent progression (January 2024), showing right pleural effusion and progressive pleural thickening consistent with malignant mesothelioma. (B) Representative bright-field images of the mesothelioma patient-derived organoid (MPM-PDO) model during early culture. Organoid structures are visible after one week, progressively developing branching extensions and interconnected cord-like networks. (C) Histologic and immunohistochemical comparison between the primary mesothelioma biopsy and the corresponding MPM-PDO model. H&E staining shows irregular and stellate morphology; immunohistochemistry demonstrates loss of BAP1 nuclear staining and preserved Calretinin (nuclear and cytoplasmic) and WT1 (nuclear) expression in both the primary tumor and organoids, confirming phenotypic concordance.

In parallel, the establishment of PDOs from the mesothelioma patient’s pleural effusion was feasible, with organoid structures observed after one week in culture (Figure 3B; left panel). In contrast to the NSCLC-PDO models, these MPM-PDOs evolved exhibiting branching extensions with cords of cells that eventually interconnected, forming a network-like structure (Figure 3B; middle and right panels). Concordantly, haematoxylin and eosin (H&E) staining revealed an irregular, stellate morphology consistent with the observed 3D growth pattern, closely resembling the desmoplastic response seen in the primary mesothelioma biopsy (Figure 3C); the magnification in the upper right corner exhibiting an atypical mitosis. Immunohistochemical profiling confirmed the expression of canonical mesothelioma markers, including loss of BAP1 nuclear staining, concordant in both the primary tumor and the MPM-PDO model (Figure 3C). Additionally, organoids showed strong Calretinin positivity with combined nuclear and cytoplasmic expression, and positive nuclear labeling of WT-1 (Figure 3C), underscoring the phenotypic fidelity of the MPM-PDOs to the original tumor tissue.

To evaluate the clinical utility of the mesothelioma preclinical model based on organoids derived from the malignant pleural effusion, we conducted a pharmacological study to first confirm that the MPM-PDO model mimicked the response to the standard therapy observed at the clinical setting, and second, to analyze the performance of the malignant mesothelial PDO model for the screening of alternative therapies that could be eventually translated into the clinical practice. To study the sensitivity of the mesothelioma PDO model to the standard therapy based on carboplatin, we performed a dose-response cell viability assay by exposing the organoids to increasing concentrations of carboplatin for 72 hours, as described previously for the NSCLC-PDO models. As shown in Figure 4A, the dose-response curve of sensitivity to carboplatin showed an IC50 of 0.23 mM, compatible with a highly resistant preclinical model. Similar to the NSCLC-PDOs, these results lead us to conclude that the malignant mesothelioma PDO model mirrors the absence of response to first line therapy shown by the patient, and confirmed this model as faithfully recapitulating the clinical characteristics of the malignant mesothelioma of origin.

**Figure 4.**
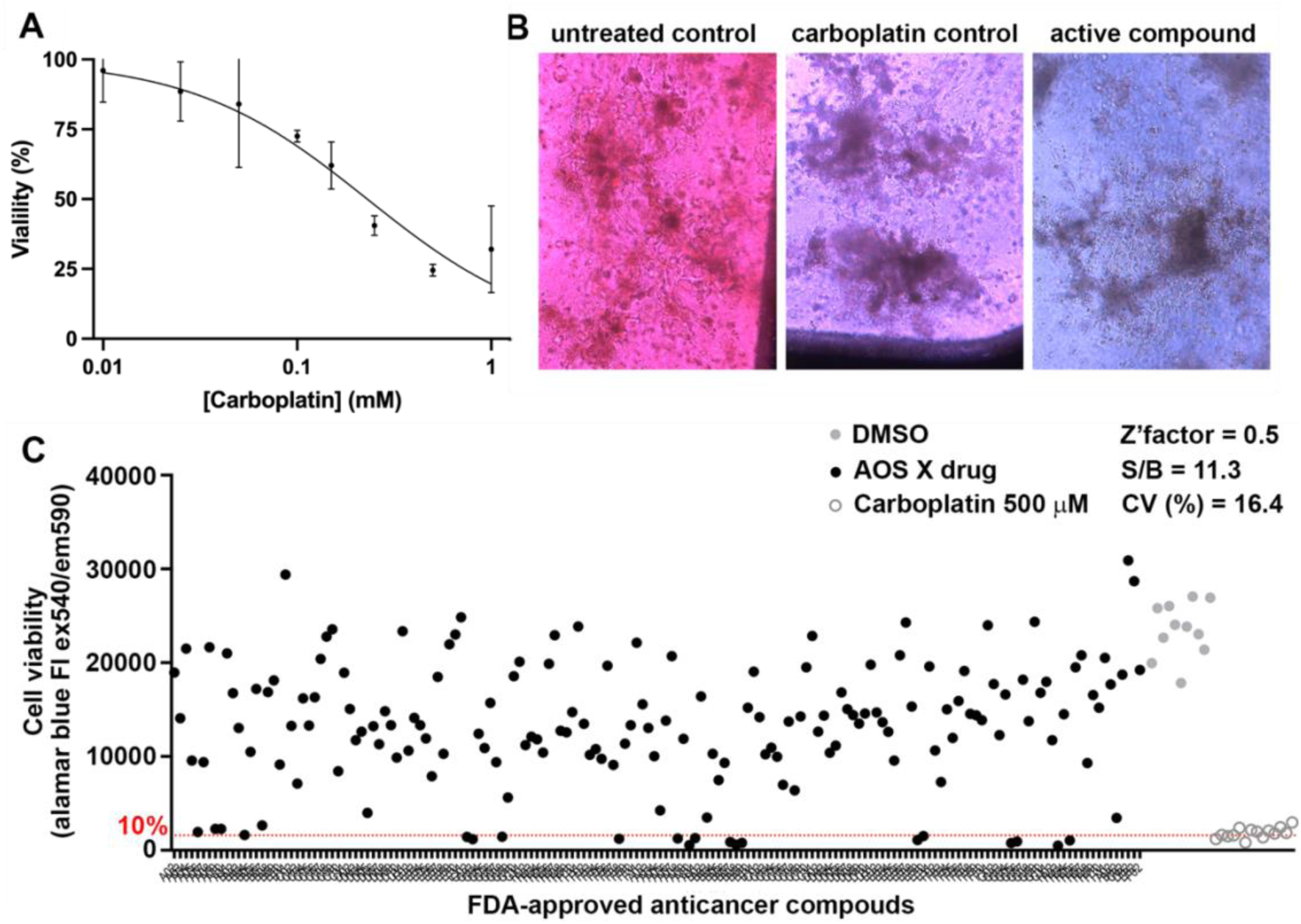
Functional validation and high-throughput drug screening of the mesothelioma PDO model. (A) Dose–response curve of the MPM-PDO model treated with increasing concentrations of carboplatin for 72 hours, showing an IC₅₀ of 0.23 mM, consistent with a resistant phenotype. (B) Representative images from the high-throughput screening assay: untreated control organoids (left), carboplatin 500 µM toxicity control (middle), and an active compound inducing marked cytotoxicity (right). (C) Scatter plot of the high-throughput screening with 169 FDA-approved anticancer agents in the MPM-PDO model. Each dot represents an individual compound; DMSO and carboplatin controls are indicated. Compounds inducing <10% viability relative to carboplatin control were considered active hits.

We finally explored the performance of the MPM-PDO model in a high throughput screening with the 169 FDA-approved anticancer agents, as previously shown for the NSCLC-PDO models. Representative images of the PDO6 colored in magenta to blue indicative of the cellular viability to cell toxicity, respectively, illustrate viable untreated organoids (Figure 4B; left panel), control of cell toxicity with carboplatin at 500 µM (Figure 4B; middle panel), and an active compound with massive apoptotic effect (Figure 4B; right panel). When referred to the mean value of the 500 µM carboplatin control, the HTS identified 16 active compounds showing <10% viability of the PDO6 (Figure 4C), notably a group of selective tyrosine kinase receptor inhibitors (TKI) highlighting the potential of this therapeutic strategy in malignant mesothelioma. The TKI selection included in addition to Vemurafenib and Tucatinib similar to NSCLC-PDOs, Abemaciclib, a dual inhibitor of cyclin-dependent kinases 4 (CDK4) and 6 (CDK6); Dabrafenib, targeting BRAF; Entrectinib and Larotrectinib, as selective inhibitors of neurotrophin receptor kinase (NTRK); and Osimertinib, a third-generation epidermal growth factor receptor (EGFR) tyrosine kinase inhibitor. Moreover, this study also identified antineoplasic antibiotics that act by targeting RNA translation and transcription, like Actinomycin-D and Plicamicyn; the anthracycline Epirubicin, and the protein synthesis inhibitor Omacetaxine mepesuccinate. Finally, the high throughput screening conducted with the MPM-PDO model identified Idarubicin, Carfilzomib, Tazemetostat, Panobinostat and Romidepsin, also with activity in the NSCLC-PDOs, suggesting potential similar alternative therapeutic strategies.

## DISCUSSION

In this work, we described the generation of patient-derived organoids from malignant pleural effusion in two thoracic neoplasias, NSCLC and mesothelioma; both share important clinical features, particularly in advanced stages, commonly involving the pleural cavity leading to the development of malignant pleural effusion, and with limited chemotherapy responses and alternative clinical options. PDO models are increasingly positioning as a preferred preclinical models that recapitulate not only the histological and molecular features of the tumors of origin, but also their structural and functional behavior making them the optimal models to explore novel pharmacological approaches.

We generated five different NSCLC-PDO models and one MPM-PDO model from pleural effusions, confirming their concordance with the tumors of origin. In the representative case, the primary tumor was described as an invasive acinar-pattern adenocarcinoma characterized by irregular glandular structures embedded in desmoplastic stroma. The corresponding PDO demonstrated well-formed three-dimensional glandular (acinar-like) structures with clear luminal differentiation. Immunohistochemically, diffuse cytoplasmic CK7 expression and strong nuclear TTF-1 positivity were retained in the organoid, supporting preservation of pulmonary adenocarcinoma differentiation, despite the absence of Napsin A expression. The moderate successful rate in the generation of PDO models from pleural effusion could be related to the amount and impact of the microenvironment and to the proliferative capabilities of the purified tumoral compartment; ongoing work upon segregation of both cellular components of the pleural effusion and co-culture at NSCLC-PDO model generation, will provide with evidences to understand their crosstalk during pleural dissemination and eventually on immunotherapy response.

Importantly, the PDOs were amenable to high-throughput screening, enabling the evaluation of a wide range of FDA-approved anticancer agents. Recent work on PDOs derived from malignant pleural effusions, demonstrated their suitability as models for precision oncology (15). The performance of this novel preclinical models in terms of response to therapy also reproduced the clinical behavior of the patients, representing an optimal preclinical tool for the research and evaluation of alternative therapies targeting those molecular events associated with this difficult-to-treat malignancies. Several studies have reported the successful establishment of NSCLC organoids from surgical specimens, core biopsies, and malignant pleural effusions, showing high concordance with the original tumors in terms of histopathology, genomic alterations, and drug response patterns (18,23,24). Importantly, NSCLC-PDOs have been increasingly applied to medium- and high-throughput drug screening platforms, enabling the functional evaluation of standard chemotherapies, targeted therapies, and rational drug combinations. In selected cases, organoid-based drug sensitivity profiles have correlated with clinical responses, highlighting their potential role in functional precision oncology.

Although Idarubicin showed clear cytotoxic activity in the NSCLC-PDO models, the observed IC₅₀ values were above the reported maximum plasma concentration (C_máx_=0.123 µM) achievable in clinical settings (25). This suggests that, despite its in vitro activity, Idarubicin may not represent a clinically feasible therapeutic option in this context. These findings highlight the importance of interpreting drug sensitivity results within pharmacokinetic constraints and prioritizing compounds that exert antitumor effects at concentrations within the therapeutically achievable range. In this regard, tyrosine kinase inhibitors such as Tucatinib, the proteasome inhibitor Carfilzomib, and the EZH2 inhibitor Tazemetostat demonstrated inhibitory activity within a range more compatible with clinically attainable exposure, supporting their potential consideration for further translational evaluation, either as monotherapy or in rational combination strategies (25). Likewise, similar conclusions have been achieved also in malignant peritoneal mesothelioma organoid models (26), and previous work also highlighted the impact of mesothelioma-associated to the response to chemotherapy (27), the value of murine malignant mesothelioma organoids to study strategies to overcome chemotherapy resistance (28), or malignant pleural mesothelioma organoids derived from tissue biopsies for on-chip chemotherapy screening (29).

These models are particularly relevant in patients without actionable mutations or with resistance to conventional treatments, where genomic profiling alone may be insufficient to guide therapeutic decision-making. To this regard, our results highlight the potential of epigenetic-based therapies in NSCLC and MPM models. Epigenetic dysregulation plays a significant role in the pathogenesis of both thoracic malignancies with alterations affecting chromatin remodeling complexes, histone methylation, and acetylation contributing to tumor progression, immune escape, and therapeutic resistance (30). In NSCLC, epigenetic-targeted strategies such as histone deacetylase (HDAC) inhibitors and EZH2 inhibitors have been explored in preclinical models and early-phase clinical trials, either as monotherapies or in combination with chemotherapy and immunotherapy. Similarly, mesothelioma frequently harbors alterations in chromatin regulatory genes, including BAP1, supporting a strong biological rationale for targeting epigenetic mechanisms (31). Although clinical benefit remains under active investigation, epigenetic modulators represent a promising therapeutic avenue in thoracic malignancies, particularly in tumors lacking actionable genomic drivers or displaying resistance to standard treatments (32,33). Also, the observed responses to tyrosine kinase inhibitors are of particular interest, since these agents are not part of the current standard of care for mesothelioma but may represent alternative therapeutic opportunities. While our findings are preliminary and limited to a single case, they support further exploration of targeted therapies in this setting.

The fact that this work is based on a limited number of malignant pleural effusion-derived PDO models represents a main limitation and the results cannot be generalized. In addition, drug responses observed in vitro may not fully reflect the complexity of the tumor microenvironment or in vivo conditions. Even so, the successful generation and screening of PDOs from pleural effusion shows that this approach is feasible and may support future research in thoracic malignancies. In summary, our study shows that PDOs derived from pleural effusion provide a feasible model for NSCLC and mesothelioma testing drug sensitivities. This approach may help identify new treatment options beyond standard strategies for patients with advanced NSCLC and MPM.

## Conclusion

We have developed pioneering patient-derived organoid models of non-small cell lung adenocarcinomas and of malignant mesothelioma from pleural effusion that faithfully recapitulated the morphology and histology of the tumors of origin. This models allowed testing of different drugs, including standard chemotherapies and targeted therapies, and identifying novel treatments that might of benefit for patient with these thoracic diseases. By providing a reliable and flexible platform, it offered a step forward toward more personalized approaches to treating this aggressive diseases.

---

**Supplementary Figure 1.**
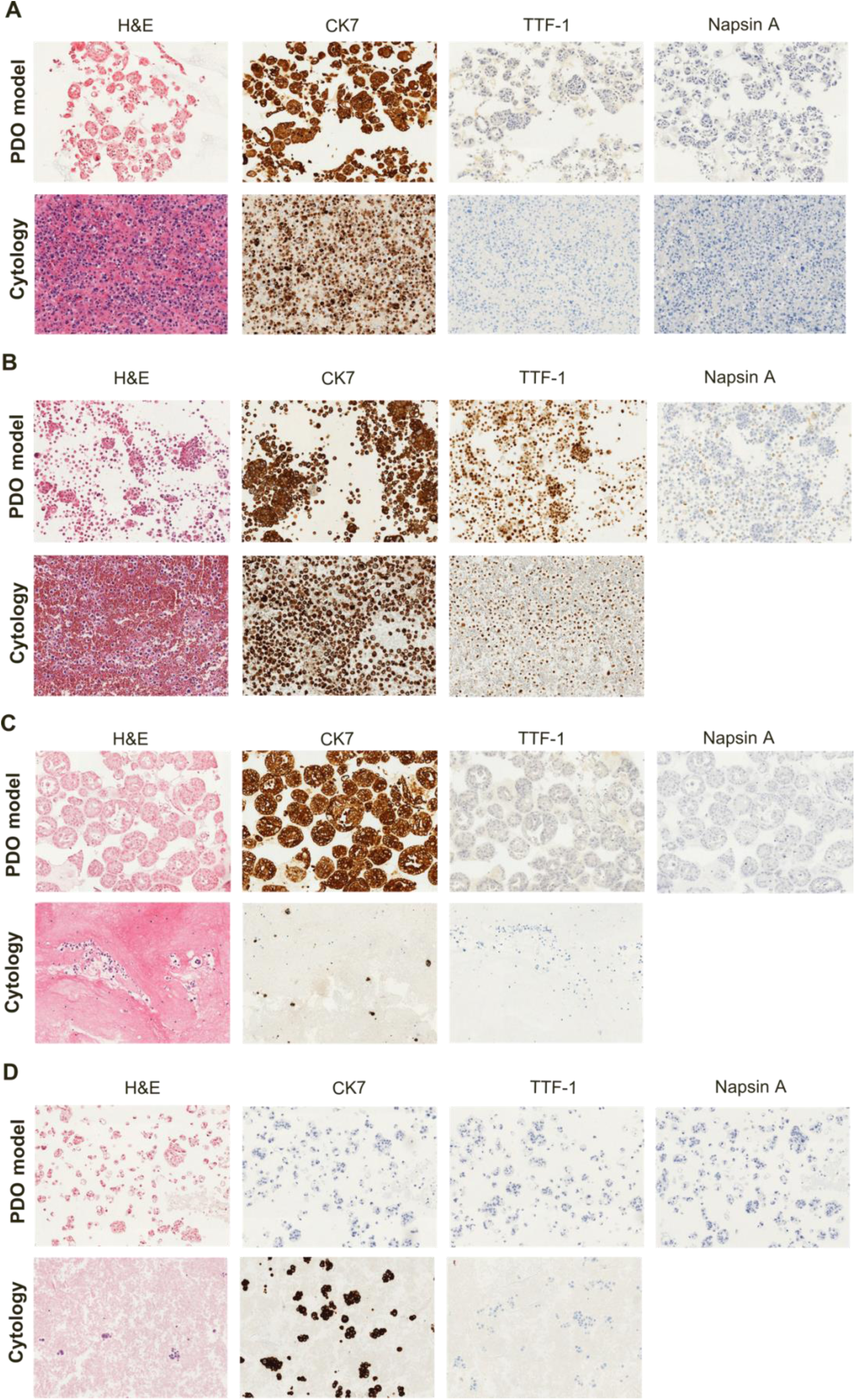
Histologic and immunophenotypic characterization of additional NSCLC pleural effusion–derived organoid models. (A–D) Representative hematoxylin and eosin (H&E) staining and immunohistochemical analysis of PDO2 (A), PDO3 (B), PDO5 (C), and PDO6 (D), compared with their corresponding cytology specimens. For each model, organoids (upper row) and matched cytology samples (lower row) are shown. Immunostaining includes CK7, TTF-1, and Napsin A. Across models, organoids retain epithelial morphology and preserve lineage-defining markers, including diffuse CK7 expression and nuclear TTF-1 positivity, consistent with pulmonary adenocarcinoma differentiation. Variability in Napsin A expression was observed among models, in line with the primary samples.

**Supplementary Figure 2.**
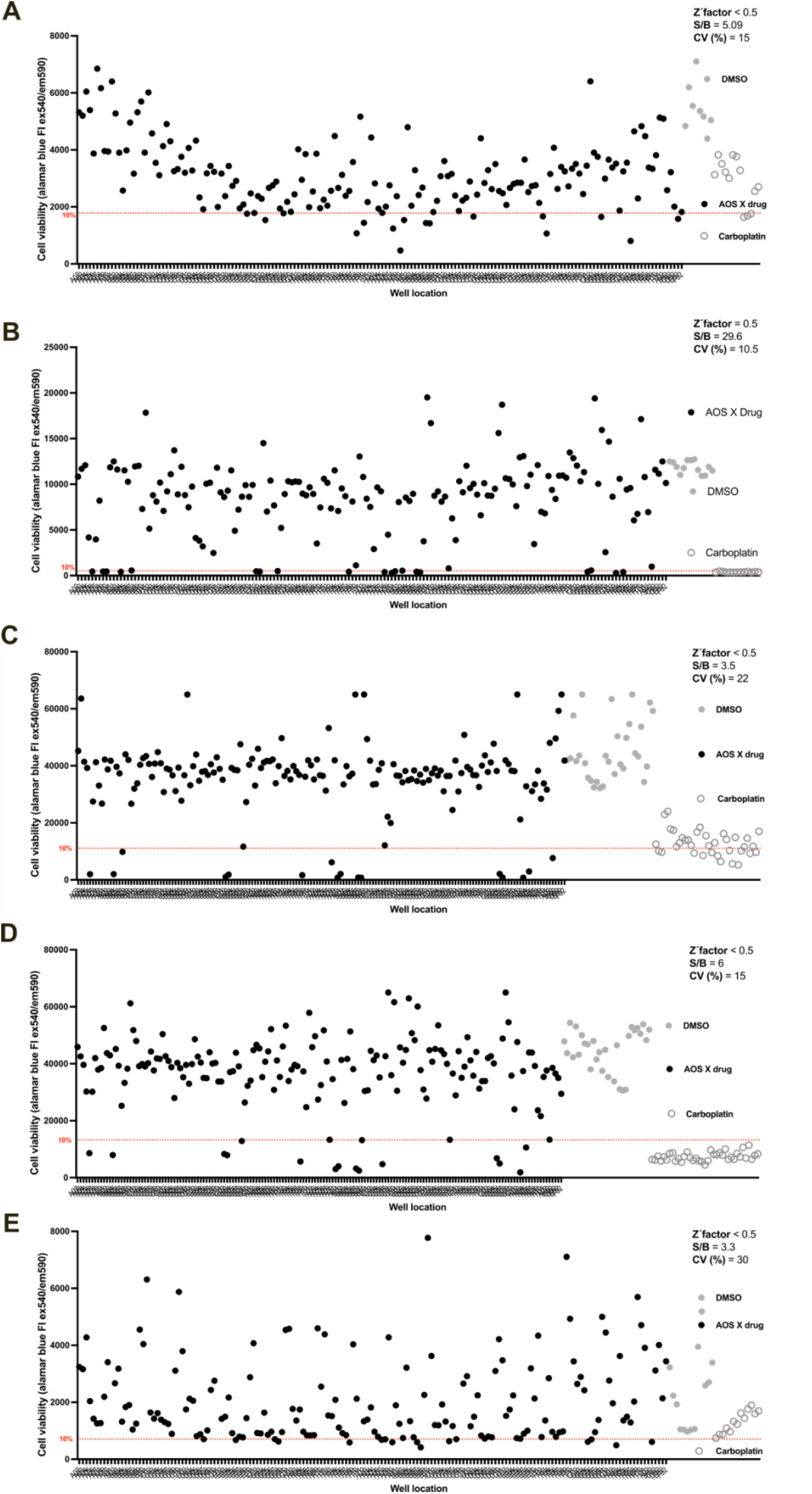
High-throughput screening scatter plots of additional NSCLC pleural effusion–derived organoid models. (A–E) Representative scatter plots of high-throughput screening with 169 FDA-approved anticancer agents in PDO2 (A), PDO3 (B), PDO5 (C), PDO5 BIS (D), and PDO6 (E). Each dot represents an individual compound tested at 10 µM for 72 hours, and cell viability was determined by AlamarBlue fluorescence (540/590 nm). DMSO-treated wells define maximal proliferation, and carboplatin 500 µM represents the toxicity control. Compounds inducing marked reduction in viability relative to controls were considered active hits.

